# Optical genome mapping identifies a novel *PSIP1::TBL1X* fusion in metastatic pancreatic neuroendocrine tumors

**DOI:** 10.64898/2025.12.17.694687

**Authors:** Daniel Ackerman, Elisabeth R. Seyferth, Wuyan Li, Julia E. Youngman, Jelsia N. Cottone, Cecilia A. Wright, Robyn T. Sussman, Abashai A. Woodard, David C. Metz, Bryson W. Katona, Michael C. Soulen, Daniel M. DePietro, Jennifer R. Eads, Terence P. Gade

## Abstract

**Background and aims:** Effective treatment of metastatic neuroendocrine tumors (NETs) is limited by a lack of targeted therapies and clinically useful predictive biomarkers. We applied complementary genomic profiling technologies to identify genomic alterations that could lead to actionable drug targets and biomarkers.

**Methods:** Optical genome mapping (OGM) and whole exome sequencing (WES) were performed on 70 liver metastases of neuroendocrine tumors from multiple anatomical primary sites. PacBio Iso-Seq long read sequencing and western blotting was used to confirm fusion gene expression. Findings were validated by rtPCR and western blotting on a separate cohort of 17 resection specimens from 14 pancreatic NET cases.

**Results:** Somatic variant calling from OGM data detected recurrent fusions involving *TBL1X* (*PSIP1::TBL1X*) and *BEND2* (*CHD7::BEND2* and *NEO1::BEND2*) in pancreatic neuroendocrine tumors (pNETs). *BEND2* fusions were associated with high grade tumors, as previously described. The novel *PSIP1::TBL1X* fusion was confirmed to result in fusion transcript expression by both long-read sequencing and nested rtPCR. Presence of the *PSIP1::TBL1X* fusion transcript was also confirmed in a second cohort of pNET resection specimens and fusion protein expression was established by western blotting. Both *TBL1X* and *BEND2* fusions are mutually exclusive with *ATRX/DAXX* mutations.

**Conclusion:** This study underscores the importance of structural variants, including fusion genes as both prognostic biomarkers and potential therapeutic targets, especially in *ATRX/DAXX* negative pNETs. *BEND2* fusion positivity may be a clinically useful biomarker for aggressive disease. As all *TBL1X* fusion isoforms contain the entire *TBL1X* coding sequence, TBL1X inhibition could be explored as a novel treatment for *TBL1X* fusion positive pNETs.

## Introduction

Neuroendocrine tumors (NET) are a rare and heterogenous cancer type that can develop at multiple anatomical sites and are most frequently found in the pancreas and the gastro-intestinal and bronchopulmonary tracts. While well differentiated NETs can follow a relatively indolent disease course, a large fraction of patients present with metastatic disease (mNET), with highly variable prognoses based on tumor grade, degree of metastasis and primary tumor location^1–3^. Despite the availability of several therapeutic options, including both systemic and locoregional therapies, treatment selection for mNET remains challenging and new biomarkers of clinically meaningful subtypes are urgently needed to target therapies more effectively^4,5^.

Short-read sequencing has been the predominant approach taken in molecular profiling studies of NETs. This technology is well suited to the identification of small alterations, such as single nucleotide variants (SNVs) and small insertions and deletions (InDels), but is less sensitive for the detection of structural variants (SVs)^6^. SVs make up a large percentage of driver mutations in many cancer types^7^ and can drive cancer progression through loss of tumor suppressors, amplified expression of oncogenes and production of novel fusion oncogenes, many of which can be successfully targeted.

Long read sequencing and optical genome mapping (OGM) platforms overcome several technical challenges of accurate and sensitive SV calling^8,9^, but show unequal sensitivities to SVs of different types^10^. OGM has been successfully applied in both clinical and research settings to understand complex chromosomal events^8,9,11^ and to identify chromosomal abnormalities associated with cancer subtypes^12^. Here we combined OGM with short read sequencing to characterize a cohort of liver metastases of NETs, identify recurring fusion genes, and further characterize a previously unreported *PSIP::TBL1X* gene fusion.

## Materials and Methods

### Ethics statement

This retrospective study was conducted following the Declaration of Helsinki and approved by the University of Pennsylvania institutional review board (IRB). Biopsy specimens were obtained under an observational clinical trial (IRB approval #825782) collecting mNET biopsies prior to locoregional therapy using ultrasound guidance. Resection specimens were obtained from the Penn Neuroendocrine Tumor Program Biobank (IRB #814805). There was no patient involvement in the design, conduct, or dissemination of this research.

### OGM sample preparation

Ultra-high molecular weight (UHMW) DNA was extracted from tumor tissue using the Bionano Prep SP Tissue and Tumor DNA Isolation Kit or from buffy coat blood samples using the SP-G2 Blood & Cell Culture DNA Isolation Kit following the relevant Bionano protocols (Document #30339 and CG-00006 revision B) with minor modifications. UHMW DNA was labeled using the DLS-G2 Labeling kit (protocol #CG-30553-1 revision E) and loaded on Saphyr Chips. OGM datasets were analyzed using the Rare Variant Analysis pipeline with reference genome build hg38, Bionano Solve v3.8.1, Bionano Access v1.8.1.

### Somatic variant calling by NGS

For WES, UHMW DNA samples were diluted to 50ng/µl in 20µl and vortexed at highest setting for 30sec prior to re-quantification by NanoDrop (ThermoFisher). Whole exome sequencing was performed by Novogene on a NovaSeq X Plus Series using 150bp paired end sequencing and yielded approximately 48Gb per sample. Somatic variant calling by GATK and variant annotation was performed using a panel of normals (PON). Targeted sequencing using the TSO500 panel (Illumina), was run by the University of Pennsylvania Genomic Analysis Core using standard protocols. To adapt UHMW DNA for input, DNA was diluted to 1ng/µl in a volume of 130µl and sheared using the Covaris LE220 in single tubes. Pooled libraries were assessed using a MiSeq Nano and run on a NovaSeq S1 flowcell.

### Identifying candidate fusions from OGM data

We extract predicted fusion cases from the PutativeGeneFusion column in smap output files and tabulated the frequency of exact fusions and of each fusion partner gene across primary subgroups. To highlight previously identified fusion genes and fusion partner genes, these data were then cross-referenced with the GDB2 gene fusion database^13^. To identify high confidence fusion cases, we considered the GDB2 degree-of-frequency (DoF) score^14^, the number of cases in our cohort containing the exact fusion, the number of cases containing each of the fusion partners in a predicted fusion, and conducted extensive literature searches.

### PacBio Iso-Seq

RNA was successfully extracted from 4 liver biopsy samples using the PureLink RNA mini kit (Thermo Fisher Scientific) and submitted for PacBio Kinnex sequencing at the University of Washington Long Read Sequencing Center, generating 9.4M HiFi reads. Long-read fusion transcript detection was performed using CTAT-LR-fusion v1.1.1^15^ executed through a Singularity container. The container image (ctat_lr_fusion.v1.1.1.simg) was obtained from the project repository (https://github.com/TrinityCTAT/CTAT-LR-fusion/wiki/ctat_lr_fusion_docker_and_singularity). Analysis used the GRCh38_gencode_v37_CTAT_lib_Mar012021.plug-n-play reference library downloaded from https://data.broadinstitute.org/Trinity/CTAT_RESOURCE_LIB/.

### Nested PCR

Both inner and outer PCR reactions were run at a final volume of 25µl using 12.5µl Phusion Hot Start Flex 2X Master Mix (NEB), primers at a final concentration of 500nM and the following cycling conditions: 98°C for 30sec, 20 or 35 cycles of (10sec 98°C, 30 sec 59.8°C, 30sec 72°C), 5 min 72°C. The first ‘outer’ reaction was run using primers PT10 (5’-AGACGAAGTTCCTGATGGAGC-3’) and PT9 (5’-GTGCAAGACTGACTGCATGG-3’) with up to 5µl of cDNA for 20 cycles. The resulting PCR product was diluted 10x and 1µl was added to a second “inner” PCR reaction using a nested primer pair (PT2 5’-CCACCCACAAACAAACTACC-3’ and PT8 5’-GTGGCAGCACGATGAAGAGG-3’) for 35 PCR cycles. Nested PCR reactions were run on cDNA generated using random priming of RNA extracted from fresh-frozen samples. ACTB amplification was performed using the same reagent set and was used as a reference gene control, with 0.1µl of the same cDNA sample, primers PT11 (5’-CACCATTGGCAATGAGCGGTTC-3’) and PT12 (5’-AGGTCTTTGCGGATGTCCACGT-3’), and the following cycling conditions: 98°C for 30sec, 20 or 35 cycles of (10sec 98°C, 30 sec 59.8°C, 30sec 72°C), 5 min 72°C. All oligos were synthesized by Integrated DNA Technologies and were dissolved in molecular grade water to a stock concentration of 100µM.

### TOPO cloning

A-overhangs were added to PCR products using 1µl of 1mM dATP, 0.3µl of 50mM MgCl2, 1µl of 10x Taq polymerase Buffer and 0.2µl (1 unit) of Taq (Invitrogen) for 15min at 72°C in a total volume of 10µl prior to cloning. The resulting product was directly cloned using the TOPO TA Cloning Kit for Sequencing (Invitrogen) following the manufacturer’s instructions. Sanger sequencing was performed by the University of Pennsylvania DNA sequencing core facility.

### Western blotting

Twenty micrograms (20µg) of protein samples, lysed in RIPA buffer containing Pierce Protease and Phosphatase Inhibitor Mini Tablets (ThermoFisher Scientific), were denatured using Bolt LDS sample buffer (Life Technologies) containing sample reducing agent (Life Technologies) and were heated to 98ºC for 5 min. Denatured samples were run on Bolt BisTris 4-12% gels and transferred to PVDF membranes using iBlot3 Transfer Stacks (Invitrogen) using broad range transfer conditions (6min, 25V, low cooling). Membranes were blocked for at least 1h at room temperature in 5% milk in TBST buffer (Cell Signaling) and blocked membranes were incubated overnight at 4ºC in 5% milk TBST containing primary Ab.

| Antibody | Cat # | Dilution | Incubation |
| --- | --- | --- | --- |
| anti- TBL1X | 13540-1-AP | 1:400 | O/N, 4°C, in 5% milk |
| anti- β-actin | 20536-1-AP | 1:10000 | O/N, 4°C, in 5% milk |
| anti- Rabbit HRP | 7074S | 1:7500 | 1h, RT, in 5% milk |

## Results

### Optical genome mapping identifies recurrent TBL1X and BEND2 fusions

OGM was performed on the Saphyr platform using UHMW DNA extracted from fresh frozen biopsy tissue of liver metastases from 70 patients with mNET. Our cohort included metastases from a variety of primary sites, including small bowel (n=23), pancreas (n=30, 28 pNET, 2 pancreatic neuroendocrine carcinoma (pNEC)), lung/bronchus (n=6), colon/rectum (n=5), four cases with unknown primary sites, and one case each of appendix and stomach primary NETs. Matched normal control samples were included for a majority of patients (n = 55) and Bionano rare variant analysis (RVA) was performed to identify somatic structural variants. SNVs and small InDels were identified by either WES or targeted sequencing using the TSO500 panel of cancer-related genes. Our data revealed several known features specific to pNETs, including frequent mutations in *ATRX, DAXX* and *MEN1*^16^ (Supp Fig 1, Supp Tables 1-3). pNETs were also found to have higher numbers of SVs overall, including frequent alterations in *PTPRD, DMD, MACROD2* as well as in the *MTAP/CDKN2A/CDKN2B* locus (Figure 1A, Supp Fig 1).

**Figure 1.**
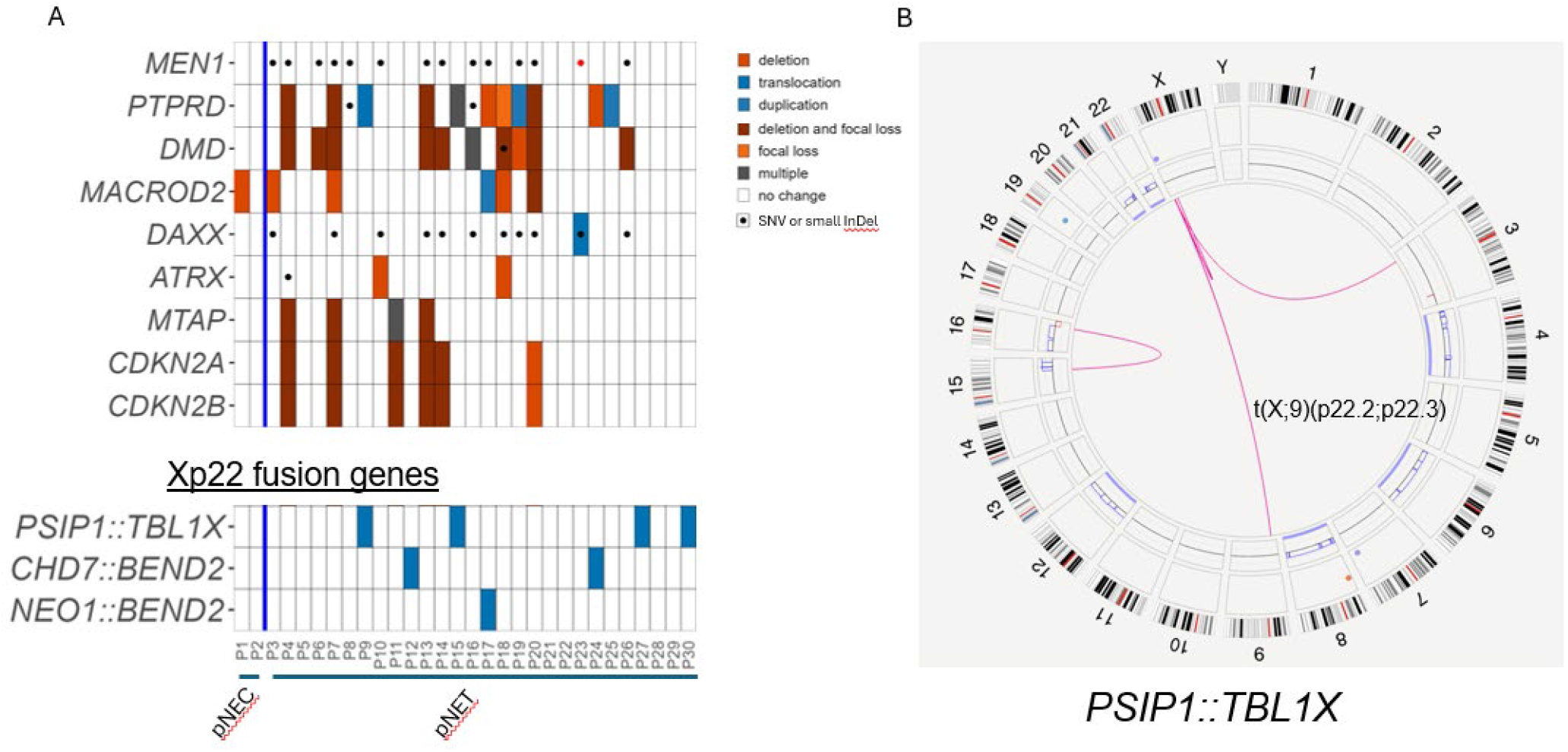
**(A)** Heatmap showing the most common alterations identified in liver metastases of pancreatic neuroendocrine tumors and carcinomas by OGM and WES. P1-P30: pancreatic NET or NEC samples. **(B)** Representative (Sample P27) circos plot showing t(X;9)(p22.2;p22.3) translocation with breakpoints in the PSIP1 and TBL1X gene regions

As expected, given the higher overall number of genomic rearrangements, the number of predicted fusion genes was also highest in pNETs (Supp Fig 2). In order to identify fusion genes that may play an oncogenic role, we examined each partner gene using the Fusion Gene annotation DataBase GDB2^13^. Several recurring fusions were found to have exact matches, had been previously characterized, or showed high degree-of-frequency (DoF) scores for one of the partner genes (see Suppl Table 4). We identified recurrent fusions involving the two genes *BEND2 (CHD7::BEND2, NEO1::BEND2)* and *TBL1X (PSIP1::TBL1X)*, both within the p22 genomic region of the X-chromosome (Figure 1B, Suppl Fig 3-4). *BEND2* has recently been reported as a recurrent fusion gene partner in pNET cases^17^, while *TBL1X* fusions appear to be novel. The *PSIP1::TBL1X* fusion was predicted as a result of a recurring t(X;9)(p22.2;p22.3) translocation (Figure 1B, Supp Fig 3) with remarkably consistent breakpoints. All 3 cases of *BEND2* fusions and 4/5 *PSIP::TBL1X* cases were found in pNETs, while a single case of *PSIP1::TBL1X* was identified in a NET of unknown primary (see Table 2). Neither fusion was identified in either of the pNEC cases included in this cohort (Figure 1A) and no other Xp22 genes formed recurrent fusions in our cohort, though individual cases of fusions involving *CCNB3* and *GAGE12H*, present elsewhere on the X chromosome p arm, were identified (Suppl Table 4). Predicted fusions between *MTAP, CDKN2A* and *CDKN2B*-*AS1* were not investigated further, as deletions in the 9p21 region are common due to presence of the *CDKN2A* tumor suppressor and are unlikely to lead to expression of a functional fusion oncogene.

**Table 1.** Table of high-confidence predicted fusions. GDB2, Fusion Gene annotation DataBase; DoF, degree of frequency score. ‘H gene’ and ‘T gene’ denote the 5’ and 3’ partner genes in each fusion.

| Predicted Fusion | Positive cases | Pancreas/<br>sm.bowel/<br>unknown | Fusion GDB2 DoF |  | Fusion partner frequency |  |
| --- | --- | --- | --- | --- | --- | --- |
|  |  |  | H Gene | T Gene | H Gene | T Gene |
| *PSIP1::TBL1X* | 5 | 4/0/1 | 1372 | 4232 | 5 | 5 |
| *MTAP::CDKN2B-AS1* | 4 | 3/1/0 | 6776 | 25 | 4 | 4 |
| *CDKN2A::CDKN2B-AS1* | 1 | 0/1/0 | 9375 | 25 | 1 | 4 |
| *CHD7::BEND2* | 2 | 2/0/0 | 0 | 0 | 2 | 3 |
| *NEO1::BEND2* | 1 | 1/0/0 | 3600 | 18 | 1 | 3 |
| *BEND2::PRRG1* | 1 | 1/0/0 | 18 | 108 | 3 | 1 |

**Table 2:**
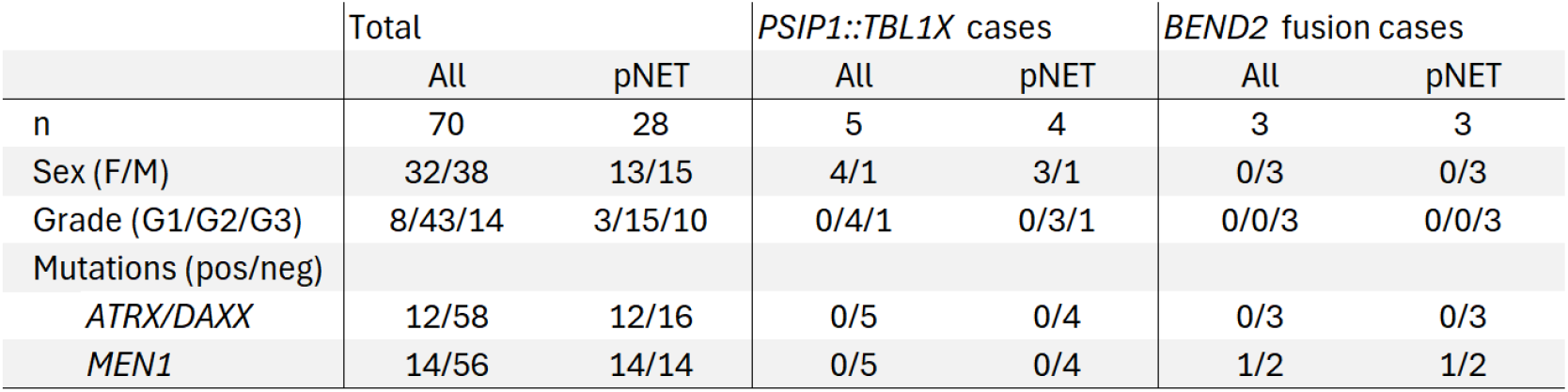
Comparison of clinical features, common gene variants in fusion positive and negative cases.

**Figure 2.**
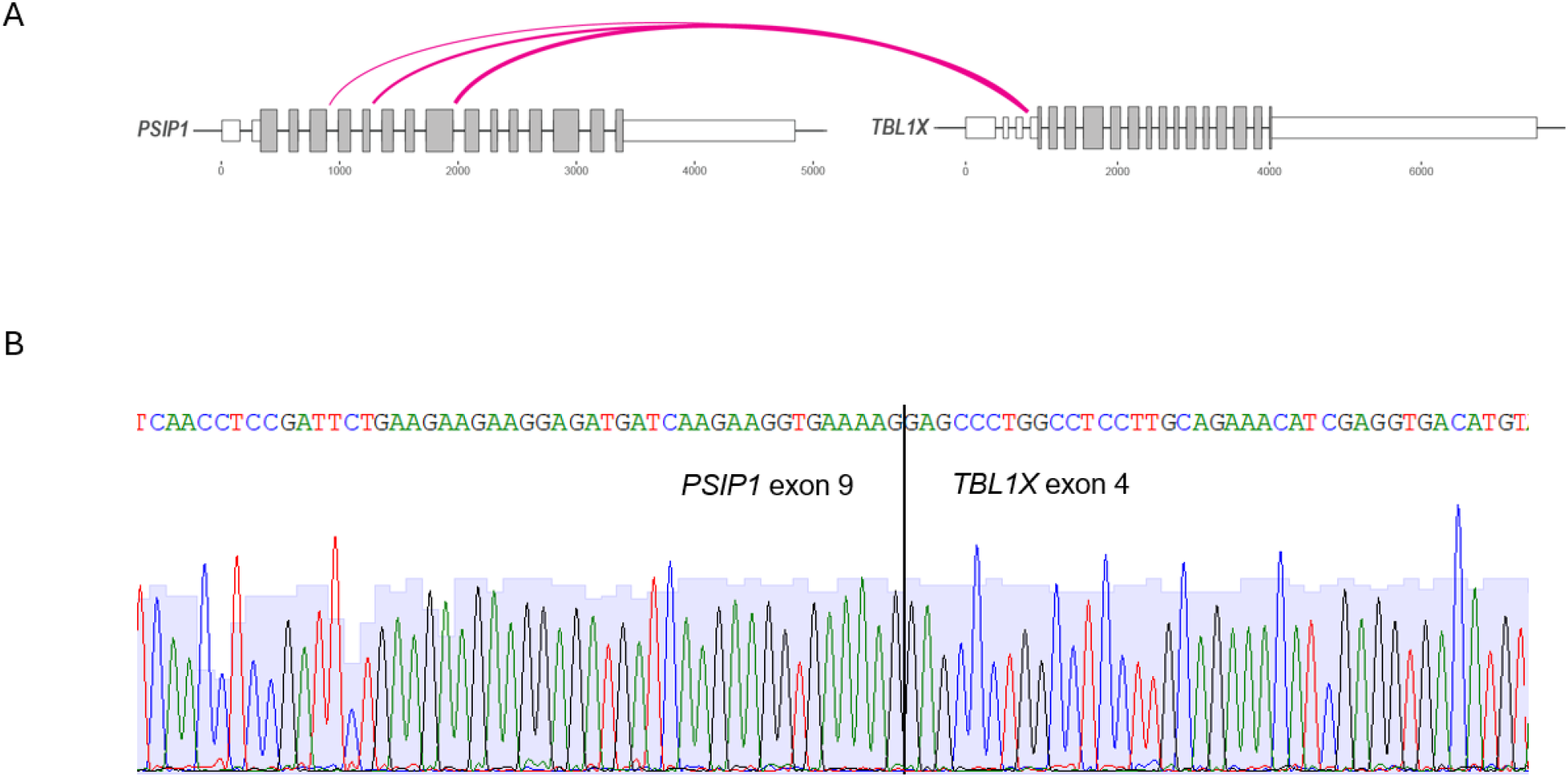
**(A)** Graphical representation of common breakpoints within *PSIP1* and *TBL1X* gene regions. Loop thickness represents frequency of the resulting isoform. **(B)** Sanger sequencing of the nested rtPCR product of *PSIP1::TBL1X(ex9-ex4)* sample P9 following TOPO cloning.

**Figure 3.**
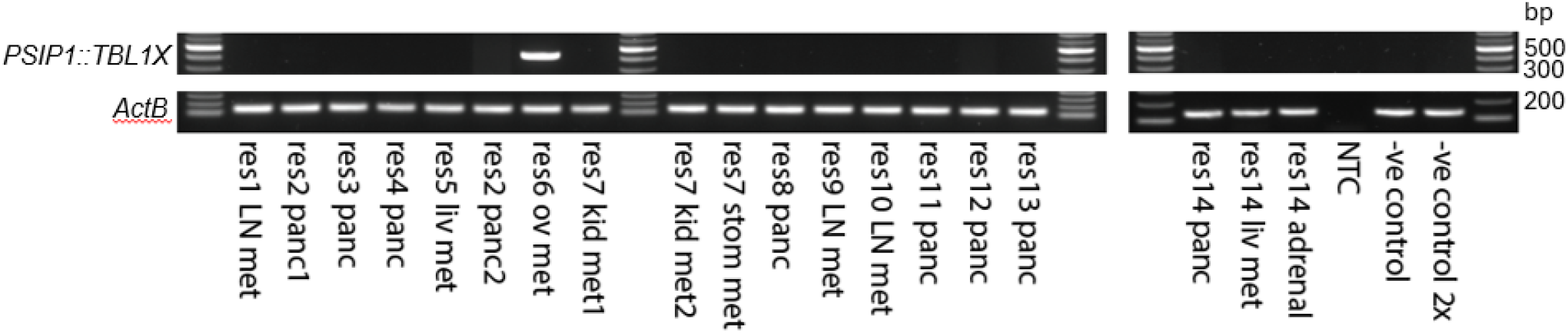
Nested rtPCR assay on resection specimen cohort showing positive *ex6-ex4* fusion variant. (res, resection specimen; LN, lymph node; met, metastasis; panc, pancreas; liv, liver; ov, ovarian; kid, kidney; stom, stomach; NTC, non-template control)

### TBL1X and BEND2 fusions are mutually exclusive with ATRX/DAXX mutations

As *BEND2* fusions have previously been found to be mutually exclusive with mutations in either the *ATRX or DAXX* gene^17^, we investigated the *ATRX/DAXX* mutation status. *ATRX/DAXX* mutations were present in 42.8% of pNET cases (Table 2) and were not found in other primary subtypes. Both *TBL1X* and *BEND2* fusion genes were mutually exclusive with *ATRX/DAXX* mutations in our cohort and both were also mutually exclusive with other common mutations, including deletions within the *DMD* and the *CDKN2A/2B/MTAP* loci. *MEN1* mutations were found in 50% of pNETs and while not mutually exclusive with BEND2 fusions, did not co-occur with *PSIP1::TBL1X* in this cohort.

Consistent with previous reports associating *BEND2* fusions with aggressive disease^17^, 3/3 *BEND2* fusions were found in grade 3 (G3) tumors and BEND2 fusions were statistically significantly associated with G3 vs G1/2 (p= 0.037 by Fisher’s exact test). *BEND2* fusions have been previously associated with female gender^17^; however, we identified BEND2 fusions exclusively in male samples (Table 2). Four out of five *PSIP1::TBL1X* cases were found in female samples, but no statistically significant association of *PSIP1::TBL1X* with gender or tumor grade was found in our cohort. pNET patients with *BEND2* fusions have been found to have heightened gastrin and ACTH secretion^17^. While none of the *BEND2* fusion cases were tested for ACTH, a patient carrying the *CHD7::BEND2* fusion showed highly elevated gastrin levels and a clinical diagnosis of Zollinger-Ellison syndrome. A second *CHD7::BEND2* positive patient had a clinical history of severe gastric ulcers but clinical plasma gastrin results were not available. These results are consistent with previous studies showing elevated gastrin expression in pNET tumors carrying BEND2 fusions^17^.

### Validating expression of *PSIP1::TBL1X* fusion transcript

While OGM is a highly sensitive method for the detection of structural variants, its resolution is insufficient to unambiguously determine the exon composition of predicted fusion transcripts. To confirm expression and characterize fusion transcript isoforms, we extracted RNA from additional biopsy tissue of 4/5 *PSIP::TBL1X* positive tumors for further molecular characterization. We confirmed expression of *PSIP::TBL1X* fusion transcripts from long read RNA sequencing in all four samples using PacBio Iso-Seq and the CTAT-LR fusion caller^15^ (see Suppl Table 5). This approach identified two different fusion variant transcripts, with *PSIP1::TBL1X(ex4-ex4)* identified in 1 sample and *PSIP1::TBL1X(ex9-ex4)* identified in 3 samples. The breakpoints predicted by OGM corresponded to the correct intronic region in 3/4 cases for *PSIP1* and 3/4 cases for *TBL1X*, supporting the need for complementary sequencing approaches to identify exact breakpoint locations following OGM. For further validation and as a screening tool, we developed a nested rtPCR (Supp Fig 5A) approach to determine the presence of the fusion transcript based on the known breakpoints. rtPCR amplification from fusion positive cases resulted in expected amplicon sizes and breakpoints identified by long read sequencing. Additionally, a third fusion transcript, *PSIP1::TBL1X(ex6-ex4)* was identified in 1 sample (patient P27), which showed both *PSIP1::TBL1X(ex6-ex4)* and *PSIP1::TBL1X(ex9-ex4)* isoforms, suggesting that the *PSIP1::TBL1X* fusion may have occurred twice in this tumor. All breakpoints were confirmed by Sanger sequencing of TOPO-cloned PCR products (Figure 2, Suppl Table 6).

### Validation of fusion in a cohort of resection specimens

An independent cohort of pNET resection samples was subsequently screened for the presence of the *PSIP1::TBL1X* fusion by nested rtPCR. These samples included 17 pNET resection specimens from 14 patients, including 9 primary tumor samples from 8 patients and 9 metastases (3 lymph node, 2 liver, 2 kidney, 1 stomach, 1 ovary) from 7 patients. No matched primary tumor and metastatic tumor samples from the same patient were available from this cohort. This approach identified an ovarian pNET metastasis sample with an amplicon matching the expected size for a *PSIP1::TBL1X(ex6-ex4)* transcript (Figure 3, Supp Fig 5B). These breakpoints were confirmed by Sanger sequencing of the TOPO cloned PCR product. We thus observed *PSIP1::TBL1X* fusion in 14% (4/28) of pNET liver biopsy samples, 0% (0/8) of primary pNET resections and 14% (1/7) of pNET metastasis resection cases. While our sample size remains relatively small, these data suggest the *PSIP1::TBL1X* fusion is more frequent in pNET metastases than primary tumors. Assuming sufficient sensitivity of the nested rtPCR assay alone and equal frequency of the *PSIP1::TBL1X* fusion in biopsied and resected metastases, we estimate a frequency of 14.3% in metastatic pNET samples (95% conf. interval: 4.8%-30.3%, Pearson-Klopper ‘exact’ binomial method).

To confirm that the presence of the in-frame fusion transcript results in expression of a fusion protein, we performed western blotting on the protein lysates from the *PSIP1::TBL1X(ex6-ex4)* positive ovarian metastasis sample and fusion negative control samples (Figure 4). As expected, this sample was found to have a TBL1X-reactive band at the predicted size (81kDa), confirming that expression of the *PSIP1::TBL1X* transcripts are translated and lead to expression of a TBL1X fusion protein.

**Figure 4.**
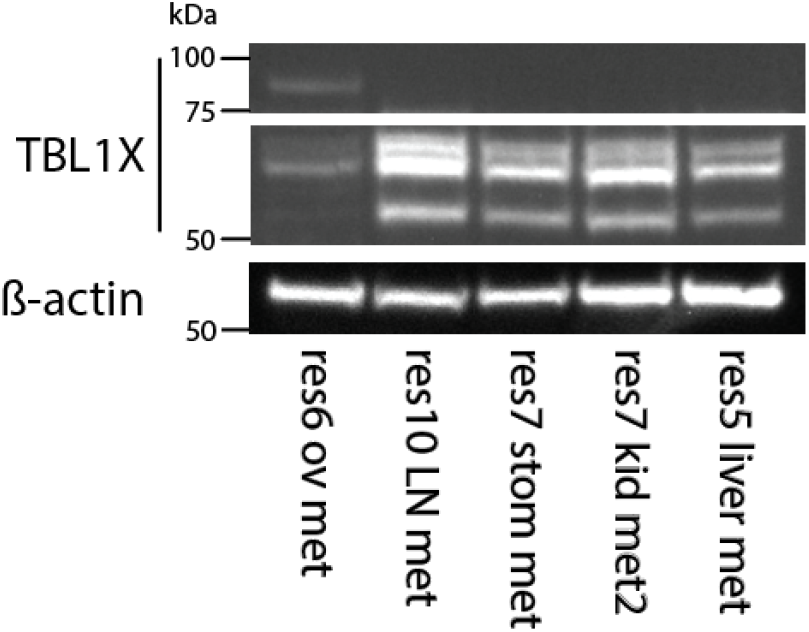
Western blot of resection specimens, including the fusion positive ‘res6 ov met’ sample (see Figure 3). PVDF membranes were probed with anti-TBL1X and anti- ß-actin antibodies.

## Discussion

We identified two genes, *BEND2* and *TBL1X* within the p22 region of the X-chromosome that form recurrent fusions with independent partner genes in pNETs. The significance of their shared genomic location is unclear, but fusions involving the X-chromosome can lead to unusual gene deregulation due to X-chromosome silencing^18^ and may also occur at a higher rate due to the elevated mutation rate found on silenced X-chromosomes^19^. Fusion genes containing *BEND2* have been described previously in several cancers^20–26^ and were recently found to be associated with aggressive disease and a unique transcriptional signature^17^ in pNETs. In our cohort, *BEND2* was fused to *CHD7* in two cases and *NEO1* in one case, but fusions to *EWSR1* have also been reported in pNETs^17,27^. Because the associated elevation in BEND2 protein levels can be detected by immunohistochemistry and is correlated with poor outcomes^17^, *BEND2* fusions are likely to be useful independent prognostic biomarkers.

The TBL1X protein is expressed ubiquitously with low tissue specificity^28^ and is involved in transcriptional gene regulation by binding the NCoR and SMRT co-repressor complexes and regulating Wnt pathway activation^29^. Its role in stabilizing ß-catenin, an important driver in several cancers, has motivated efforts to target *TBL1X* for cancer therapy^30,31^. The small molecule tegavivint is currently being tested as a TBL1X inhibitor in several ß-catenin positive cancers^32–34^. Tegavivint is believed to act by binding a hydrophobic pocket in the N-terminal region of TBL1X, impairing its interaction with ß-catenin. The exon structure of *TBL1X* fusion transcripts consistently includes the early exons of *PSIP1* fused in frame with the entire coding sequence of *TBL1X*. TBL1X fusion proteins are therefore likely to preserve native TBL1X activity and to remain sensitive to TBL1X inhibition.

The domain architecture of the *TBL1X* fusion cases we identified suggests that the PSIP1 N-terminal PWWP domain confers important functions to the fusion protein. PSIP1, also known as LEDGF, is an epigenetic reader that associates with actively transcribed chromatin and binds H3K36me3^35,36^. Beyond its role in transcriptional regulation, its functions further include regulation of splicing^37^, as well as promotion of genome integrity^38^ and DNA repair^39^. PSIP1 can form an oncogenic fusion protein with NUP98 in AML^40^ and is found to be overexpressed in several solid tumors^41^. The PWWP domain mediates chromatin binding as well as interaction with multiple protein binding partners^42^. Fusion of this domain with TBL1X may alter the localization of ß-catenin bound to the TBL1X moiety, its recruitment to chromatin or influence the proximity of PWWP binding proteins with TBL1X-containing protein complexes.

In our cohort, *BEND2* and *PSIP1::TBL1X* fusions did not co-occur with *ATRX/DAXX* mutations or common structural variants like *DMD* and 9p21 deletions. This mutual exclusivity with *ATRX/DAXX* mutations has been previously reported for *BEND2* fusions^17,43^. *ATRX/DAXX* loss is associated with genomic instability^44^ and alternative lengthening of telomeres (ALT), which may result in distinct pathogenic mechanisms and different patterns of structural variation. Our study supports the existence of a molecular pNET subtype characterized by fusion oncogenes resulting from Xp22 alterations, which is distinct from the *ATRX/DAXX* mutant subgroup.

While the *PSIP1::TBL1X* fusion has not been reported previously, careful examination of fusion database GDB2 identified a single case of *PSIP1::TBL1X(ex6-ex4)* in a pheochromocytoma sample^45^, a rare subtype of NET not included in our study. The absence of this fusion in any other cancers despite extensive RNA sequencing performed on a wide range of tumor types indicates that this fusion is highly specific for pNET. Further investigation of the functional consequences of the *PSIP1::TBL1X* fusion event may yield important insight into pNET biology.

Our study therefore opens a new avenue for the exploration of oncogenic mechanisms in NET and identifies *TBL1X* as a promising therapeutic target. Our data also support previous findings regarding the potential of BEND2 as a predictive biomarker of aggressive disease. Taken together, these findings demonstrate the promise of applying complementary technologies to improve molecular profiling and identify novel tumor biology.

## References

1. Dasari, A. et al. Epidemiology of Neuroendocrine Neoplasms in the US. JAMA Netw Open 8, e2515798 (2025).

2. Hallet, J. et al. Exploring the rising incidence of neuroendocrine tumors: A population-based analysis of epidemiology, metastatic presentation, and outcomes. Cancer 121, 589–597 (2015).

3. Hofland, J., Kaltsas, G. & de Herder, W. W. Advances in the Diagnosis and Management of Well-Differentiated Neuroendocrine Neoplasms. Endocrine Reviews 41, 371–403 (2020).

4. Couvelard, A. et al. Heterogeneity of tumor prognostic markers: a reproducibility study applied to liver metastases of pancreatic endocrine tumors. Mod Pathol 22, 273–281 (2009).

5. Shi, C. et al. Liver Metastases of Small Intestine Neuroendocrine Tumors: Ki-67 Heterogeneity and World Health Organization Grade Discordance With Primary Tumors. American Journal of Clinical Pathology 143, 398–404 (2015).

6. Dremsek, P. et al. Optical Genome Mapping in Routine Human Genetic Diagnostics—Its Advantages and Limitations. Genes 12, 1958 (2021).

7. Cosenza, M. R., Rodriguez-Martin, B. & Korbel, J. O. Structural Variation in Cancer: Role, Prevalence, and Mechanisms. Annual Review of Genomics and Human Genetics 23, 123–152 (2022).

8. Chan, E. K. F. et al. Optical mapping reveals a higher level of genomic architecture of chained fusions in cancer. Genome Res. 28, 726–738 (2018).

9. Luebeck, J. et al. AmpliconReconstructor integrates NGS and optical mapping to resolve the complex structures of focal amplifications. Nat Commun 11, 4374 (2020).

10. Savara, J., Novosád, T., Gajdoš, P. & Kriegová, E. Comparison of structural variants detected by optical mapping with long-read next-generation sequencing. Bioinformatics 37, 3398–3404 (2021).

11. Dixon, J. R. et al. Integrative detection and analysis of structural variation in cancer genomes. Nat Genet 50, 1388–1398 (2018).

12. Kriegova, E. et al. Whole-genome optical mapping of bone-marrow myeloma cells reveals association of extramedullary multiple myeloma with chromosome 1 abnormalities. Sci Rep 11, 14671 (2021).

13. Kim, P. et al. FusionGDB 2.0: fusion gene annotation updates aided by deep learning. Nucleic Acids Res 50, D1221–D1230 (2022).

14. Kim, P., Jia, P. & Zhao, Z. Kinase impact assessment in the landscape of fusion genes that retain kinase domains: a pan-cancer study. Brief Bioinform 19, 450–460 (2018).

15. Qin, Q. et al. CTAT-LR-fusion: accurate fusion transcript identification from long and short read isoform sequencing at bulk or single cell resolution. bioRxiv 2024.02.24.581862 (2024) doi:10.1101/2024.02.24.581862.

16. Gujarathi, R. et al. MEN1/DAXX/ATRX mutations enhance progression-free survival in gastroenteropancreatic neuroendocrine tumors treated with peptide receptor radionuclide therapy. Endocrine-Related Cancer 31, (2024).

17. Wood-Trageser, M. A. et al. Recurrent BEND2 Fusion Genes Identified by Whole Transcriptome Sequencing of Nonfunctional Pancreatic Neuroendocrine Tumors Correlate With Poor Patient Prognosis. Modern Pathology 38, 100863 (2025).

18. Spatz, A., Borg, C. & Feunteun, J. X-Chromosome Genetics and Human Cancer. Nat Rev Cancer 4, 617–629 (2004).

19. Jäger, N. et al. Hypermutation of the Inactive X Chromosome Is a Frequent Event in Cancer. Cell 155, 567–581 (2013).

20. Yoshida, A. et al. Soft-tissue sarcoma with MN1-BEND2 fusion: A case report and comparison with astroblastoma. Genes, Chromosomes and Cancer 61, 427–431 (2022).

21. Lehman, N. L. et al. Astroblastomas exhibit radial glia stem cell lineages and differential expression of imprinted and X-inactivation escape genes. Nat Commun 13, 2083 (2022).

22. Lucas, C.-H. G. et al. EWSR1-BEND2 fusion defines an epigenetically distinct subtype of astroblastoma. Acta Neuropathol 143, 109–113 (2022).

23. Sturm, D. et al. New Brain Tumor Entities Emerge from Molecular Classification of CNS-PNETs. Cell 164, 1060–1072 (2016).

24. Palsgrove, D. N., Manucha, V., Park, J. Y. & Bishop, J. A. A Low-grade Sinonasal Sarcoma Harboring EWSR1::BEND2: Expanding the Differential Diagnosis of Sinonasal Spindle Cell Neoplasms. Head and Neck Pathol 17, 571–575 (2023).

25. Liu, X. et al. RNA Sequencing Reveals Novel Oncogenic Fusions and Depicts Detailed Fusion Transcripts of FN1-FGFR1 in Phosphaturic Mesenchymal Tumors. Modern Pathology 36, 100266 (2023).

26. Dashti, N. K., Matcuk, G., Agaimy, A., Saoud, C. & Antonescu, C. R. Malignant Bone-Forming Neoplasm With NIPBL::BEND2 Fusion. Genes, Chromosomes and Cancer 63, e70015 (2024).

27. Scarpa, A. et al. Whole-genome landscape of pancreatic neuroendocrine tumours. Nature 543, 65–71 (2017).

28. Pray, B. A. TBL1X in Aggressive B-cell Lymphomas: Characterizing the Oncogenic Function and Potential as a Therapeutic Target. (The Ohio State University, 2024).

29. Pray, B. A., Youssef, Y. & Alinari, L. TBL1X: At the crossroads of transcriptional and posttranscriptional regulation. Experimental Hematology 116, 18–25 (2022).

30. Yang, R., Pray, B., Alinari, L., Li, P. K. & Cheng, X. Design, Synthesis, and Biological Evaluation of Selective TBL1X Degraders. ACS Med. Chem. Lett. 15, 1699–1707 (2024).

31. Soldi, R. et al. The Small Molecule BC-2059 Inhibits Wingless/Integrated (Wnt)-Dependent Gene Transcription in Cancer through Disruption of the Transducin β-Like 1-β-Catenin Protein Complex. The Journal of Pharmacology and Experimental Therapeutics 378, 77–86 (2021).

32. Cranmer, L. D. et al. Results of a phase I dose escalation and expansion study of tegavivint (BC2059), a first-in-class TBL1 inhibitor for patients with progressive, unresectable desmoid tumor. JCO 40, 11523–11523 (2022).

33. Whittle, S. B. et al. Abstract CT090: PEPN2011: a phase 1/2 study of tegavivint in children, adolescents, and young adults with recurrent or refractory solid tumors, including lymphomas and desmoid tumors: a report from the pediatric early phase clinical trials network. Cancer Res 83, CT090 (2023).

34. Li, D. et al. A phase 1/2 study of the TBL1 inhibitor, tegavivint (BC2059), in patients (pts) with advanced hepatocellular carcinoma (aHCC) with β-catenin activating mutations. JCO 42, TPS4192–TPS4192 (2024).

35. Eidahl, J. O. et al. Structural basis for high-affinity binding of LEDGF PWWP to mononucleosomes. Nucleic Acids Res 41, 3924–3936 (2013).

36. Akele, M., Iervolino, M., Van Belle, S., Christ, F. & Debyser, Z. Role of LEDGF/p75 (PSIP1) in oncogenesis. Insights in molecular mechanism and therapeutic potential. Biochimica et Biophysica Acta (BBA) - Reviews on Cancer 1880, 189248 (2025).

37. Pradeepa, M. M., Sutherland, H. G., Ule, J., Grimes, G. R. & Bickmore, W. A. Psip1/Ledgf p52 Binds Methylated Histone H3K36 and Splicing Factors and Contributes to the Regulation of Alternative Splicing. PLOS Genetics 8, e1002717 (2012).

38. Jayakumar, S. et al. PSIP1/LEDGF reduces R-loops at transcription sites to maintain genome integrity. Nat Commun 15, 361 (2024).

39. Daugaard, M. et al. LEDGF (p75) promotes DNA-end resection and homologous recombination. Nat Struct Mol Biol 19, 803–810 (2012).

40. Hussey, D. J., Moore, S., Nicola, M. & Dobrovic, A. Fusion of the NUP98 gene with the LEDGF/p52 gene defines a recurrent acute myeloid leukemia translocation. BMC Genet 2, 20 (2001).

41. Zhang, Y. et al. Identification of the H3K36me3 reader LEDGF/p75 in the pancancer landscape and functional exploration in clear cell renal cell carcinoma. Computational and Structural Biotechnology Journal 21, 4134–4148 (2023).

42. Morchikh, M. et al. TOX4 and NOVA1 Proteins Are Partners of the LEDGF PWWP Domain and Affect HIV-1 Replication. PLOS ONE 8, e81217 (2013).

43. Agaimy, A. et al. Gene fusions are frequent in ACTH-secreting neuroendocrine neoplasms of the pancreas, but not in their non-pancreatic counterparts. Virchows Arch 482, 507–516 (2023).

44. Clatterbuck Soper, S. F. & Meltzer, P. S. ATRX/DAXX: Guarding the Genome against the Hazards of ALT. Genes 14, 790 (2023).

45. Fishbein, L. et al. Comprehensive Molecular Characterization of Pheochromocytoma and Paraganglioma. Cancer Cell 31, 181–193 (2017).

