## Supplemental Figures for "Optical genome mapping identifies a novel *PSIP1::TBL1X* fusion in metastatic pancreatic neuroendocrine tumors"

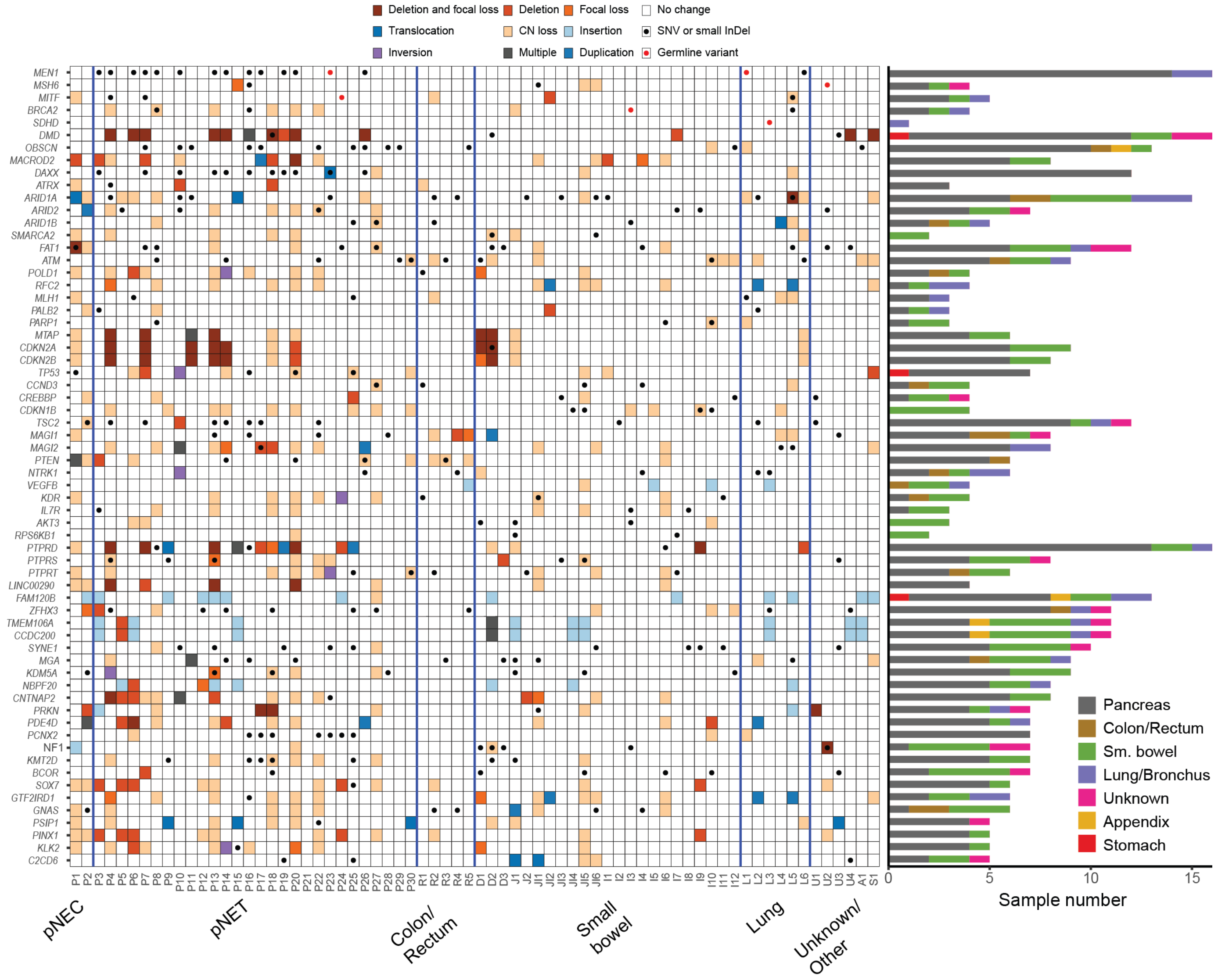


**Supplemental Figure 1:** Heatmap showing the most common alterations identified in liver metastases of pancreatic neuroendocrine tumors and carcinomas by OGM and WES. Sample identification: P, pancreas; R, colon/rectum; D, duodenum; I, ileum; J, jejunum; JI, jejunum/ileum; SB, small bowel; L, lung; U, unknown; A, appendix; S, stomach

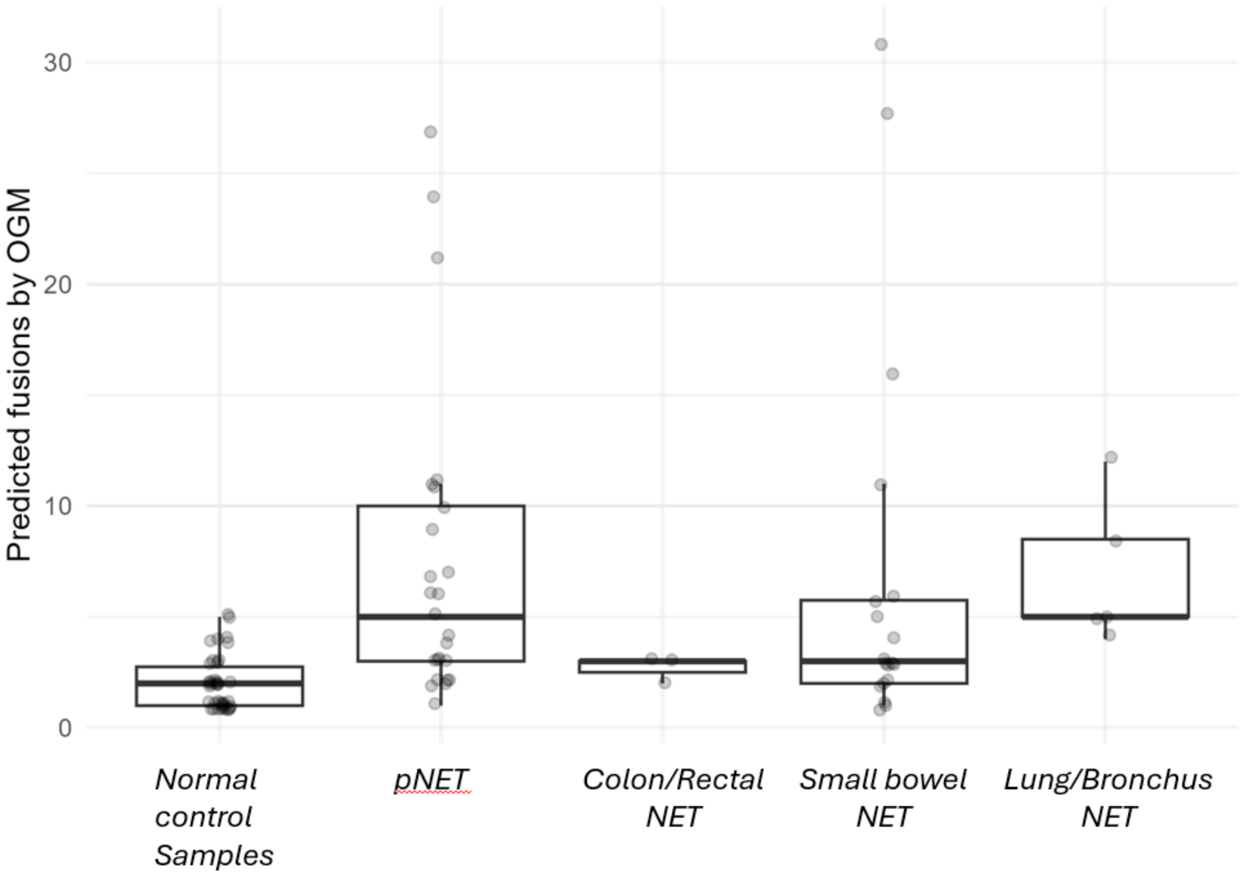


**Supplemental Figure 2:** Number of unique predicted fusions per sample in RVA analysis of OGM data across primary subtypes. All data collected from biopsy tissue of liver metastases. NEC cases were excluded (n=3).

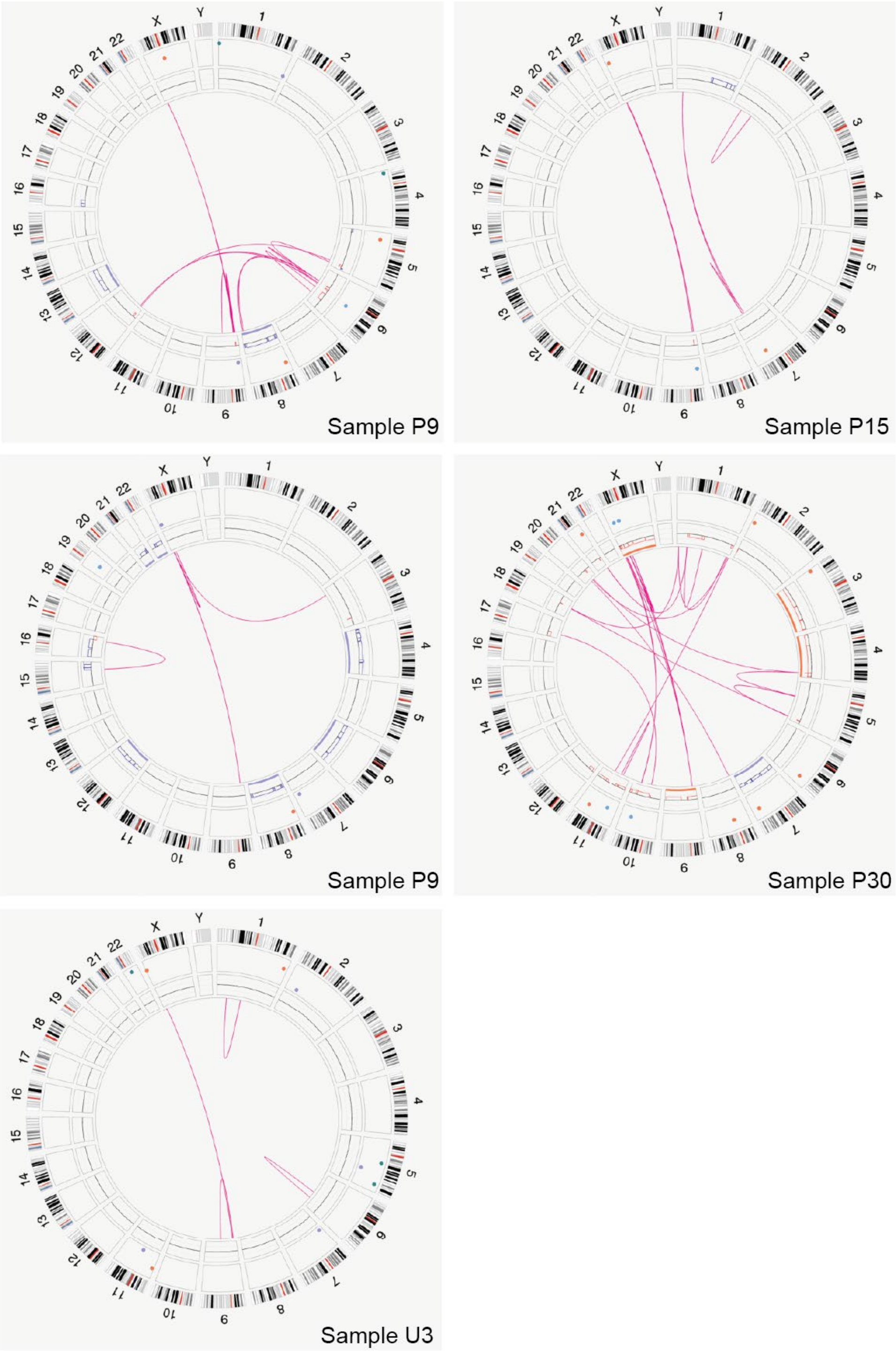


**Supplemental Figure 3:** Circos plots showing structural variants detected by OGM for samples with PSIP1::TBL1X fusion resulting from t(X;9)(p22.2;p22.3) translocation.

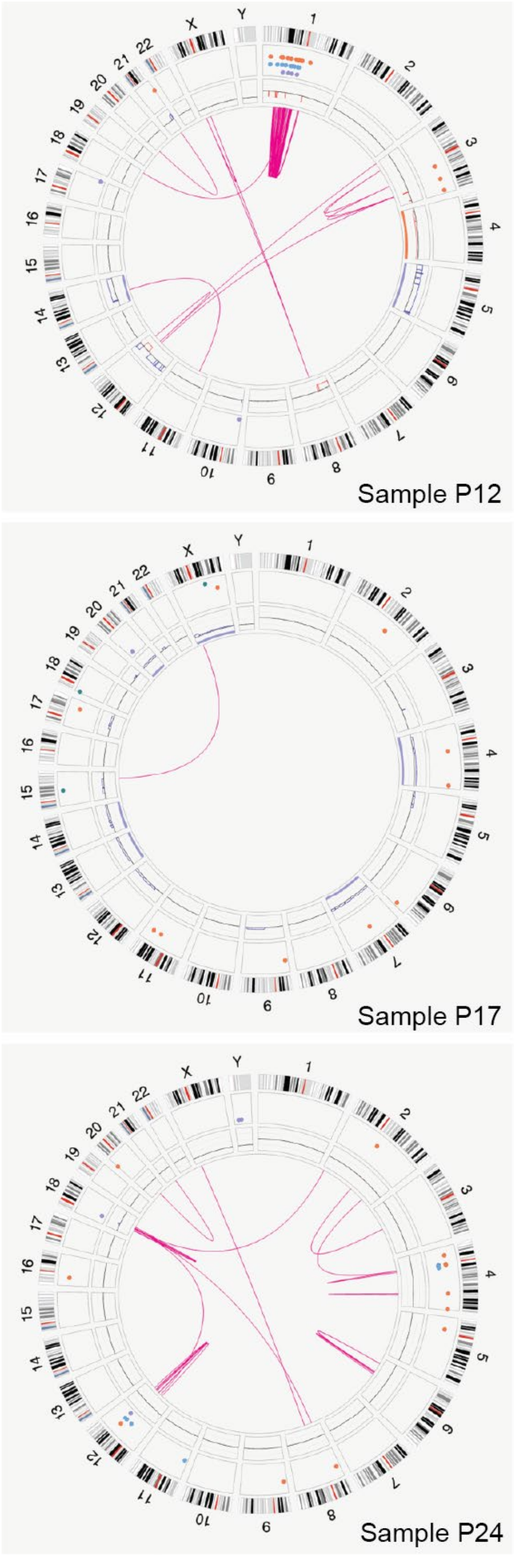


*CHD7::BEND2*
*BEND2::PRRG*

*NEO1::BEND2*

*CHD7::BEND2*

**Supplemental Figure 4:** Circos plots showing structural variants detected by OGM for samples with *BEND2* fusions

A

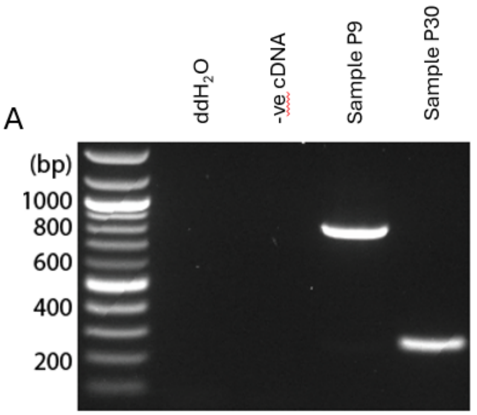


B

| Fusion isoform | Expected size |
|---|---|
| *PSIP1::TBL1X(ex4-ex4)* | 267bp |
| *PSIP1::TBL1X(ex6-ex4)* | 435bp |
| *PSIP1::TBL1X(ex9-ex4)* | 837bp |

**Supplemental Figure 5: (A)** Nested PT10/9-2/8 rtPCR assay for PSIP1::TBL1X fusion on samples P9 (*ex9-ex4* fusion variant) and P30 (*ex4-ex4* fusion variant) showing corresponding amplicon sizes. **(B)** Expected PT10/9-2/8 rtPCR amplicon sizes for each fusion isoform
